# Mutant huntingtin expression in the hypothalamus promotes ventral striatal neuropathology

**DOI:** 10.1101/2023.03.04.530949

**Authors:** Rana Soylu-Kucharz, Natalie Adlesic, Marcus Davidsson, Tomas Björklund, Maria Björkqvist, Åsa Petersén

## Abstract

Huntington’s disease is a fatal neurodegenerative disorder caused by an expanded CAG triplet repeat in the huntingtin (HTT) gene. Previous research focused on neuropathology in the striatum and its association with a typical movement disorder. Direct effects of mutant HTT (mHTT) in the striatum may cause neuropathology, although non-cell autonomous effects have also been suggested. Important non-motor features of HD include psychiatric symptoms and metabolic dysfunction, which may be linked to hypothalamic neuropathology. As hypothalamic neurons project to the ventral striatum, we hypothesized that expression of mHTT in the hypothalamus leads to disrupted neurotransmission in the ventral striatum and causes pathology. The overall aim of this study was to investigate the impact of mHTT expression in the hypothalamus on ventral striatal neuropathology and its contribution to non-HD motor symptoms. We demonstrate that selective expression of mHTT in the hypothalamus leads to the loss of dopamine and cAMP-regulated phosphoprotein (DARPP-32) immunopositive neurons in the ventral striatum in mice. Contrary to the effects of direct expression of mHTT in the hypothalamus, selective overexpression of mHTT in the ventral striatum does not affect body weight. Selective expression of mHTT in the ventral striatum leads to mHTT inclusion formation and loss of DARPP-32 neurons without affecting motor activity or anxiety-like behavior. We show that DARPP-32 neuron loss in the ventral striatum is recapitulated in the R6/2 mouse model of HD. Chemogenetic activation of hypothalamic neurons projecting to the ventral striatum had a blunted response in the R6/2 mice compared to wild-type mice, indicating a disrupted hypothalamus-ventral striatal circuitry. In summary, the expression of mHTT in the hypothalamus may impact the development of ventral striatal pathology in mice. This opens the possibility that non-cell-autonomous effects in the reward circuitry play a role in HD.

## Introduction

Huntington’s disease (HD) is a fatal neurodegenerative disorder without a disease-modifying therapy [1]. An expanded CAG repeat in the huntingtin (HTT) gene causes HD, but the disease mechanisms are not fully understood [2]. While dorsal striatum pathology is related to the typical HD motor symptoms, less is known about the underlying pathology of earlier non-motor manifestations of the disease. These include reduced cognitive function, apathy, altered emotional processing, psychiatric symptoms, and metabolic dysfunction [3]. Ubiquitously expressed mutant HTT (mHTT) exerts both cell-autonomous and non-cell-autonomous effects [4]. In the basal ganglia’s most affected structure, the striatum, both cell- and non-cell autonomous effects of mHTT have been demonstrated [4]. The known non-cell-autonomous effect of mHTT in the dorsal striatum involves cortico-striatal dysfunction. The transport of brain-derived neurotrophic factor (BDNF) from cortical layers to the striatal neurons of HD is reduced due to affected vesicle transport and, consequently, triggers neurodegenerative mechanisms, including excitotoxicity and reduced striatal neurotrophic support [5-10].

The ventral striatum is one of the severely affected brain regions in the HD with a reduced number of cells and atrophy [11-13]. The ventral striatum, with the nucleus accumbens as the main component, is a major part of the limbic-motor loop and modulates reward behavior, motivation, and metabolism [14]. As part of this circuitry, the ventral striatum receives projections from the hypothalamus, including excitatory input from lateral hypothalamic neurons, including hypocretin (orexin) cells [15, 16]. The hypothalamus and the hypocretin system are affected in the clinical HD [17-20]. We previously showed that hypothalamus-specific expression of mHTT using adeno-associated viral (AAV) vectors recapitulates clinical HD hypothalamic pathology with hypocretin cell loss in mice [21]. Since hypothalamic neurons project to the ventral striatum and the HD hypothalamus is affected even before the onset of the typical motor symptoms, we hypothesized that expression of mHTT in the hypothalamus could potentially impair the ventral striatum in a non-cell-autonomous fashion. To test our hypothesis, we first assessed the effect of ventral striatum pathology in wild-type (WT) mice expressing HTT fragments selectively in the hypothalamus. We showed that long-term expression of mHTT in the hypothalamus leads to ventral striatal DARPP-32 positive cell loss without direct expression of mHTT in the ventral striatum. To assess the hypothalamus to ventral striatum neurocircuitry function in HD, we utilized the R6/2 animal model, which ubiquitously expresses the first exon of the mHTT gene and recapitulates HD clinical hypothalamic neuropathology [22-24]. We showed that DARPP-32 positive cells in the ventral striatum were reduced in the R6/2 animal model compared to WT controls. To assess the impact of mHTT expression on the hypothalamus to ventral striatum circuitry function, we employed Cre-dependent chemogenetics (Designer Receptors Exclusively Activated by a Designer Drug (DREADDs)) and retrogradely transportable Cre-expressing AAV vectors for targeting the neurocircuitry in R6/2 and WT littermates [25-27]. Chemogenetic activation of the hypothalamus to the ventral striatum neurocircuitry revealed a blunted response in R6/2 mice compared to WT controls and deteriorated rotarod performance in WT mice. Our results suggest that pathology in the hypothalamus, as one of the affected brain regions in the early HD stages, may contribute to ventral striatum degeneration in a non-cell-autonomous manner.

## Results

### Selective hypothalamic overexpression of mHTT leads to ventral striatum neuropathology

We used several AAV vectors to assess whether hypothalamus-specific mHTT expression contributes to ventral striatum pathology. First, we unilaterally injected WT mice with AAV5-GFP (green fluorescent protein) vectors in the hypothalamus and assessed GFP maker gene expression in both hypothalamus and ventral striatum at 4 weeks post-injection. Immunohistochemical labeling for GFP revealed intense GFP immunopositive hypothalamic projections to the ventral striatum (Fig. 1A). We previously showed that AAV-vector mediated hypothalamic expression of mHTT fragments recapitulates the HD clinical hypothalamic neuropathology with loss of hypocretin and reduced gene expression levels of several neuropeptides [21, 28-30]. We then used AAV-vectors expressing the first 853 amino acids (aa) of HTT injected selectively in the hypothalamus and determined effects 8 weeks post-injection (from here on noted mutant HTT fragment: HTT853-79Q and wild type HTT fragment HTT853-18Q). We assessed *HTT* expression in both the hypothalamus and the ventral striatum using quantitative RT-PCR. As mRNAs are known to be present throughout the cell, including in the axons [31], we found that HTT mRNA was also present in the ventral striatum and expressed at low levels in comparison to the significant hypothalamic HTT overexpression (Fig. 1B). We then tested whether hypothalamus-specific mHTT expression can cause ventral striatal neuropathology in WT mice. Using stereology, we quantified the number of DARPP-32 and NeuN-positive neurons in the ventral striatal area 1 year after hypothalamic injections of the AAV-vectors expressing HTT853-79Q and HTT853-18Q. DARPP-32 is selectively expressed by the medium spiny neurons of the striatum, and medium spiny neurons are the primary neurons affected in the HD brain [32]. Here, we show a 44% loss of DARPP-32 positive neurons (Fig. 1C and 1D) and a 55% loss of NeuN neurons (Fig. 1G and 1H) in the ventral striatum 1 year after injection of AAV-vectors in the hypothalamus. The reduced number of cells was not due to reduced ventral striatum area as it was comparable among groups (Fig 1E and 1H). In a parallel set of experiments, we expressed shorter fragments of HTT, using AAV-vectors expressing the first 171 N-terminal HTT with 18 or 79 polyglutamine repeats in the hypothalamus of mice. The shorter N-terminal HTT is known to be more toxic [33], and we previously showed that expression of the HTT171-79Q fragment in the hypothalamus also recapitulates HTT853-79Q-induced metabolic disturbances [21]. In the ventral striatum, the number of DARPP-32 immunopositive cells was reduced by 36% and 41% in HTT171-18Q and HTT171-79Q groups compared to uninjected controls, respectively, at 8 months post-injection (Sup. Fig. 1A and 1B). The reduction in NeuN immunopositive cell numbers was ∼20% in both HTT171-18Q and HTT171-79Q groups (Sup. Fig. 1C and 1D). We further quantified the number of cells in the ventral striatum using cresyl violet staining to assess whether the reduction in DARPP-32 and NeuN-positive cells was due to diminished transcript levels or cell loss. Assessment of the number of neurons and glial cells further confirmed cell loss in the ventral striatum without any prominent gliosis (Sup. Fig. 1E–1G). The expression of short fragments of both HTT171-18Q and HTT171-79Q in the hypothalamus resulted in hypocretin neuropathology (Sup. Fig. 1H and 1I). The body weight measurement over 32 weeks showed that hypothalamic expression of both mutant and WT short HTT fragments impacted the body weight gain comparably (Sup. Fig. 1J).

**Figure 1.**
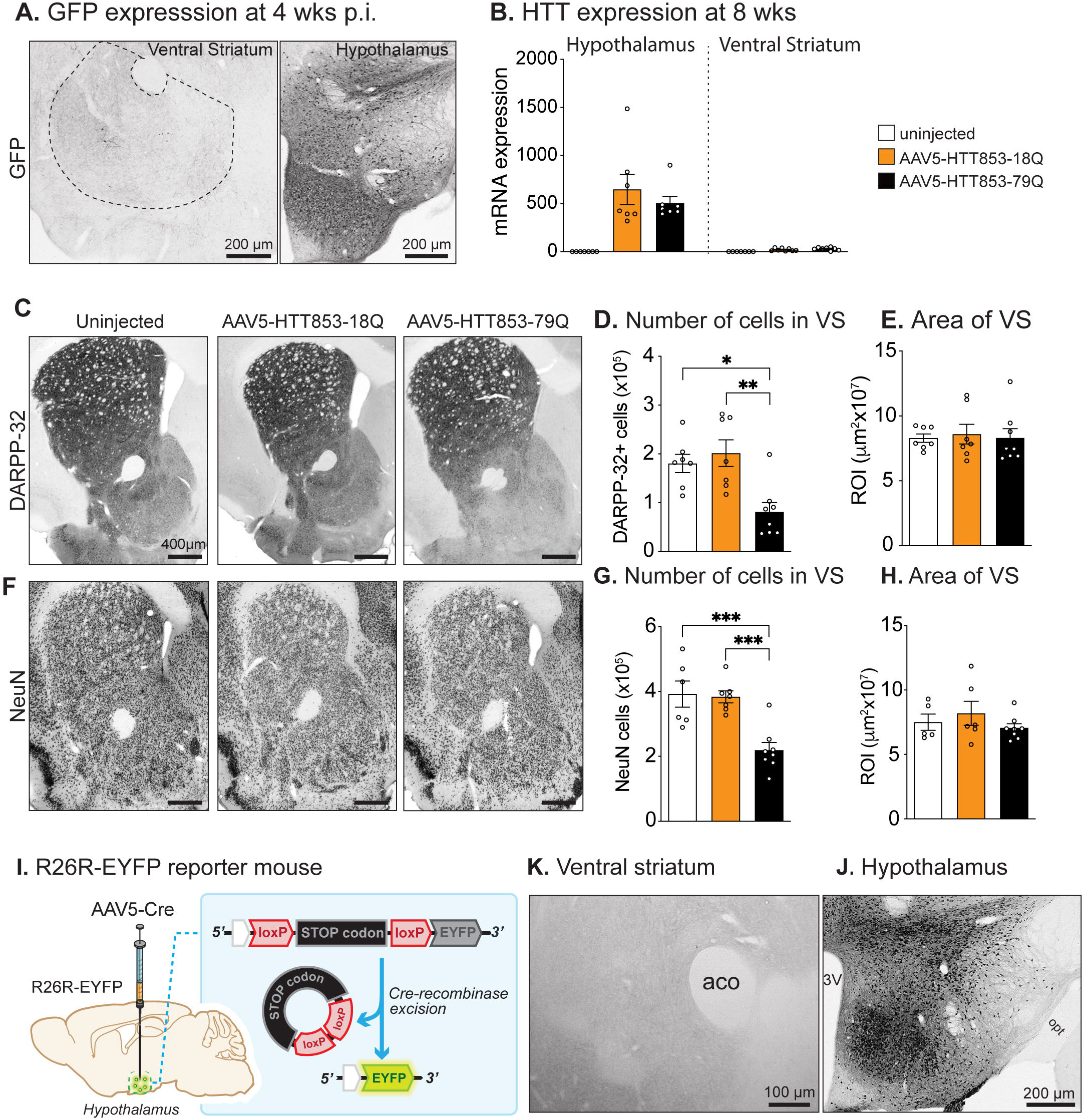
Selective mHTT expression in the hypothalamus results in ventral striatum cell loss without transgene expression in the cell soma of the ventral striatum. **(A)** Hypothalamic GFP expression at 4 weeks post-injection shows GFP-positive fibers in the ventral striatum and cell bodies in the hypothalamus. **(B)** Quantification of *HTT* mRNA levels in the hypothalamus and ventral striatum 8 weeks after the injection of AAV5-853HTT-18Q and −79Q in the hypothalamus. **(C)** Representative images show DARPP-32 immunohistochemistry in the striatum. **(D)** Stereological assessment of ventral striatum DARPP-32 positive cells shows a reduction in the AAV5-HTT-79Q group compared to both 18Q and uninjected controls (one-way ANOVA, p=0.0018). **(E)** The area of the ventral striatum was not affected in any groups as assessed via stereological analysis (one-way ANOVA, p=0.93). **(F)** NeuN immunolabeling shows striatal cells in all groups. **(G)** The number of NeuN positive cells was also significantly reduced in the 79Q group compared to 18Q and uninjected controls (one-way ANOVA, p=0.0003), **(H)** with no change in the area assessed (one-way ANOVA, p=0.43). **(I)** Illustration of Cre-dependent reporter eYFP transgene expression in R26R-EYFP mice. Expression of AAV5-Cre in the hypothalamus is validated by using rAAV5-Cre delivery in the hypothalamus of an R26R-EYFP mouse. Activation of the eYFP transcript by Cre-recombinase excision. **(J)** GFP immunohistochemistry shows the expression of GFP (EYFP) in the hypothalamic cell bodies and **(K)** absence of GFP+ cells in the striatum confirms no anterograde transport of hypothalamic transgene expression. Points on scatter graphs represent individual mice, the bars are means, and the whiskers indicate ± SEM.

Since it is known that different AAV vector serotypes can retrogradely and anterogradely propagate or passively diffuse, we next aimed to exclude the possibility of direct HTT expression in the ventral striatal neurons after hypothalamic injections of the viral vector [34]. To this end, we used rAAV5-Cre delivery in the hypothalamus of R26R-EYFP Cre-reporter mice [35]. R26R-EYFP mice harbor a floxed STOP sequence, and Cre-recombinase-mediated STOP sequence excision allows marker gene expression (Fig. 1J). Injections of R26R-EYFP mice with rAAV5 vectors that express Cre into the hypothalamus using the exact coordinates, vector volume, and titers as the HTT injection. We assessed the EYFP expression by immunobiological labeling against GFP and showed that only hypothalamic cell bodies are targeted and not cell bodies in the ventral striatal neurons (Fig. 1J and 1K and Sup. Fig. 1K). This confirms that the pathological effects in the ventral striatum are not due to the direct expression of mHTT in the striatal cells but are secondary effects of hypothalamic neuropathology.

### Ventral striatum-specific HTT expression does not affect body weight, motor activity, or anxiety-like behavior

We previously showed that hypothalamic mHTT expression in WT mice leads to hyperphagic obesity [21, 28], and inactivation of mutant mHTT selectively in the BACHD hypothalamus prevents the development of a metabolic phenotype [21, 36]. Feeding behavior and metabolic balance depend on the hypothalamic regulation [37], and hypothalamic neurons exert these functions also via reward-related circuitries to other brain regions, i.e., to the ventral striatum [16, 38, 39]. The potential metabolic effects of selective mHTT expression in the ventral striatum region have not been investigated. Since hypothalamic neuropathology led to neuronal loss in the ventral striatal area and the ventral striatum is involved in reward-related metabolic aspects, we next examined the role of HTT in this region. To this end, we delivered AAV vectors expressing GFP, HTT853-18Q, or HTT853-79Q in the ventral striatum of WT mice. First, we assessed the behavioral effects of ventral striatal HTT expression in a group of mice at 2.5 months post-injection. The body weight was comparable among the groups (Supp. Fig. 2A), and there was no effect on anxiety-like behavior as time spent in open arms and open arms entry frequency were similar among uninjected, GFP, and HTT groups in the EPM (Supp. Fig. 2B–2D). The motor phenotype was also unaltered among the groups as rotarod performance and general ambulatory movement were comparable (Supp. Fig. 2E and 2F). The expression of AAV vectors was stable for 10 months and covered the whole ventral striatal area (Fig. 2A). Expression of mHTT led to intracellular mHTT inclusion formation at 10 months post-injection (Fig. 2B). Even though mHTT expression in the ventral striatum led to 41% loss of DARPP-32 positive cells, the body weight of mice expressing HTT853-18Q and HTT853-79Q fragments was comparable at 10 months post-injection (Fig. 2D and 2E). Our data indicate that loss of ventral striatal neurons is not responsible for the development of a metabolic phenotype in mice and that the previously reported increased food intake and metabolic imbalance (Hult et al., 2011) originated from the hypothalamus specific HTT expression.

**Figure 2.**
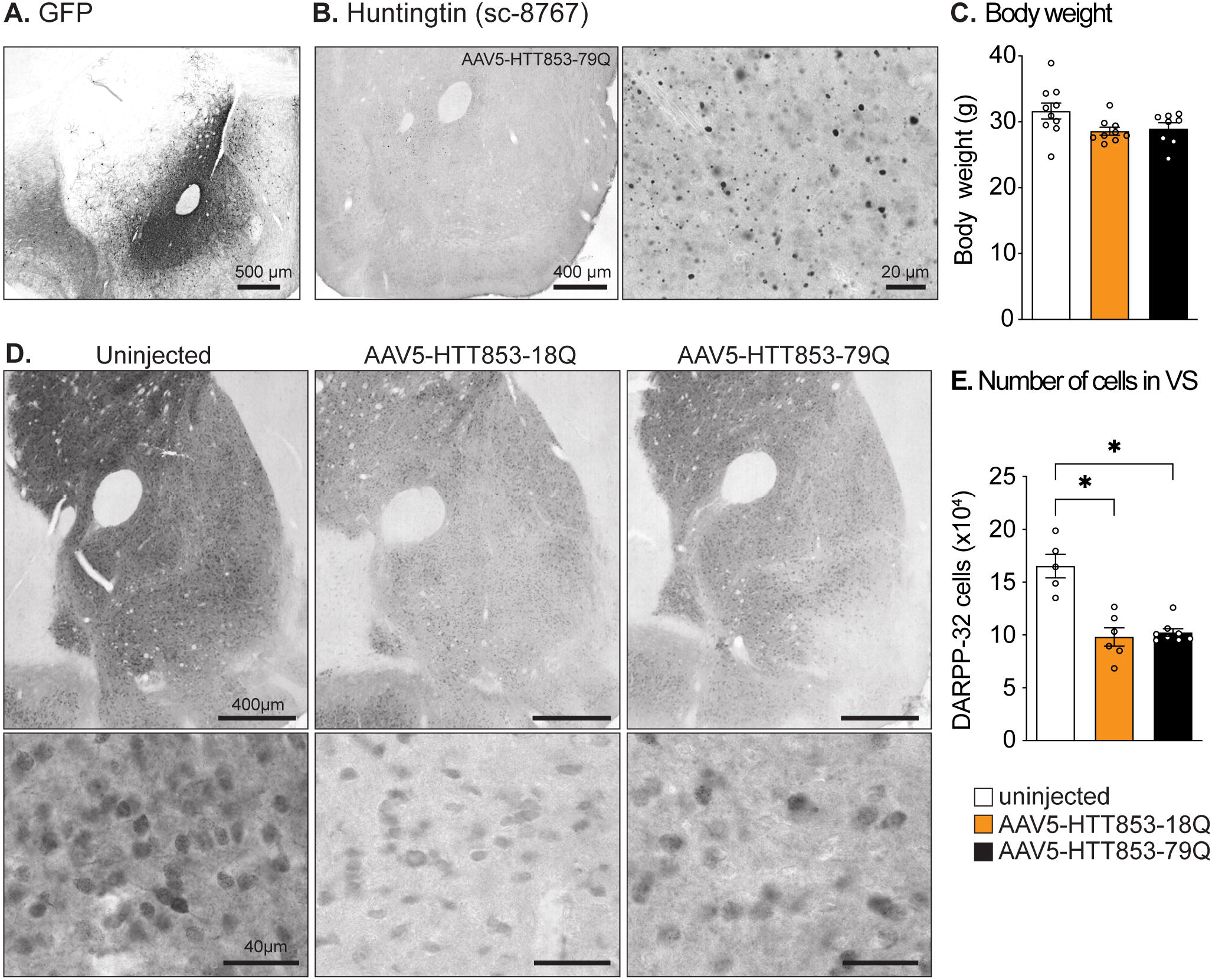
Expression of mHTT selectively in the ventral striatum of mice leads to DARPP-32 cell loss in the ventral striatum with no effect on body weight gain. **(A)** GFP and **(B)** mHTT expression in the ventral striatum at 10 months post-injection. **(C)** The body weight measurement of mice at 10 months post-injection (one-way ANOVA, p=0.059). **(D)** Representative images show DARPP-32 immunohistochemically processed sections from uninjected, HTT853-18Q, and HTT853-79Q groups. **(E)** Stereological quantification of DARPP-32 immunopositive cells in the ventral striatum shows a reduction in the number of DARPP-32 cells in both WT and mHTT expressing groups compared to uninjected mice (Kruskal-Wallis test, p=0.0013). Points on scatter graphs represent individual mice, the bars are means, and the whiskers indicate ± SEM.

### Targeting and activation of the hypothalamus to ventral striatum neurocircuitry in the R6/2 mouse model

To elucidate the function of specific neuronal populations, chemogenetic technologies have been extensively used for the last decade [40]. The combination of DREADDs with the recently introduced retrogradely transportable AAV vectors revolutionized the field of neurocircuit targeting and assessing circuit function in freely moving animals [27, 41]. To investigate the hypothalamus to ventral striatum neurocircuitry function in an HD animal model, we employed AAV vectors that express activating DREADDs and retrogradely transportable Cre-recombinase (Cre). First, we executed proof of concept experiments. To visualize hypothalamic neurons projecting to the ventral striatum, we injected retrograde AAV2-MNM004-Cre in the ventral striatum and a flex-switch color-changing AAV vector in the hypothalamus [27] (Fig. 3A and 3B). The retrograde transport of the AAV2-MNM004-Cre vector resulted in Cre-mediated excision of mCherry in the soma of hypothalamic projection neurons and allowed GFP expression only in hypothalamic neurons projecting to the ventral striatum (Fig. 3D). The rest of the hypothalamic cells not projecting to the ventral striatum expressed mCherry (Fig. 3D). Majority of the GFP-labeled cells were localized in the lateral hypothalamus, and a small population of cells was in the mediobasal hypothalamus. Using confocal imaging, we identified that most of the GFP-expressing cells were hypocretin immunopositive in the lateral hypothalamus (Supp. Fig. 3A), whereas GFP-expressing cells were not positive for oxytocin, vasopressin, or tyrosine hydroxylase in zona incerta (A13) (Supp. Fig. 3B–3E).

**Figure 3.**
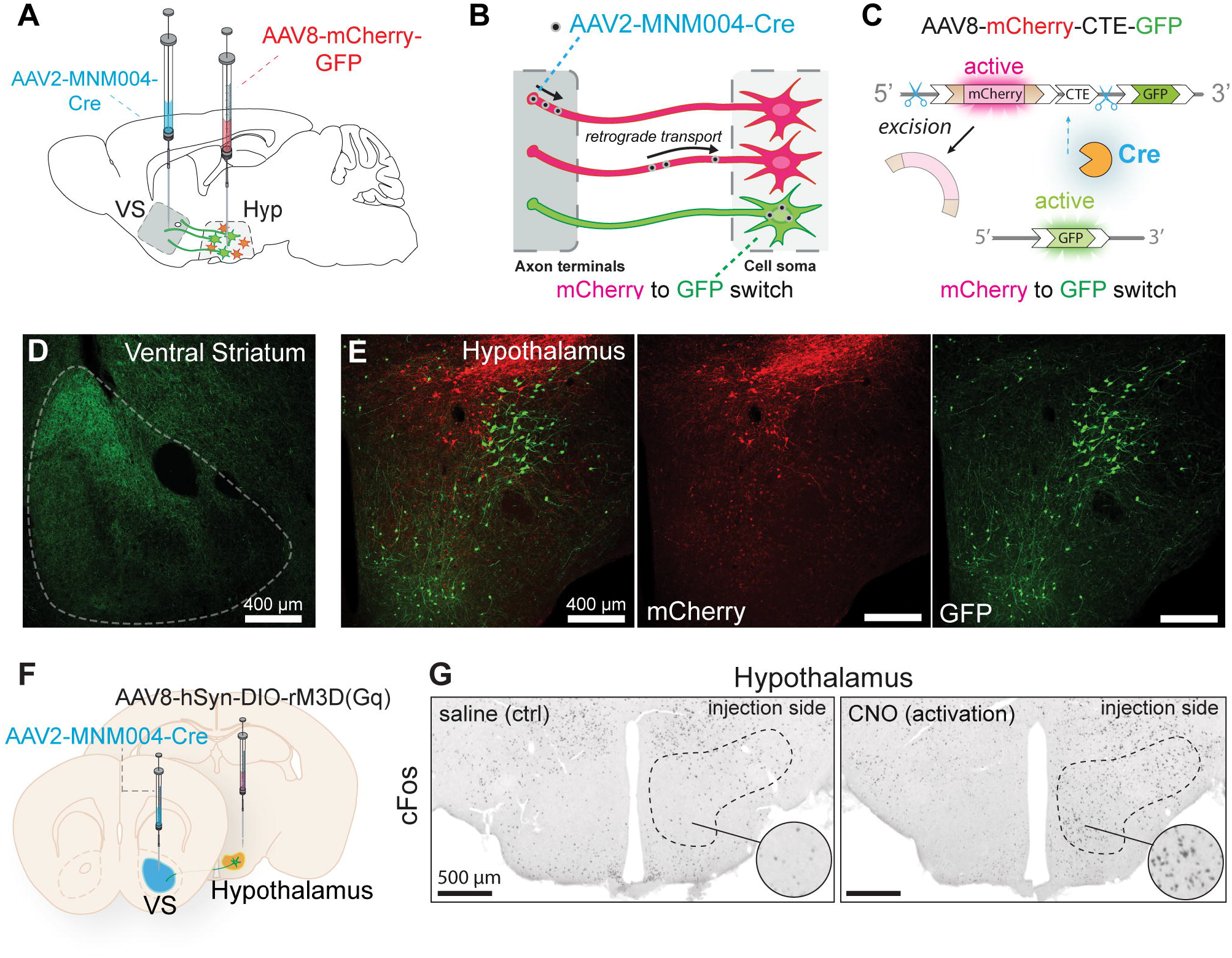
The principle of retrograde fluorescent labeling and successful activation of the hypothalamus to ventral striatum neurocircuitry. **(A)** The Cre-recombinase enzyme-expressing vector is picked up at axonal endings at **(B)** the ventral striatum injection site and transferred to cell soma in the hypothalamus, which expresses a reporter gene delivered by another viral vector, i.e., rAAV8-mCherry-CTE-GFP. **(C)** AAV2-retrograde-Cre silences mCherry in the hypothalamus and facilitates the expression of previously inactive GFP. **(D)** Cells that project from the hypothalamus to the ventral striatum express GFP, whereas other hypothalamus cells remain labeled in red by mCherry. **(E)** The principle of retrograde Cre from ventral striatum induced expression of DREADD-encoding (pAAV-hSyn-DIO-hM3D(Gq)-mCherry) exclusively in hypothalamic neurons that project to the ventral striatum. **(F)** Administration of CNO (DREADDs ligand) leads to an increase in cFos-positive cells, only in hypothalamic neurons that are part of the hypothalamus to ventral striatum signaling pathway.

Subsequently, we employed the same principle to express the Cre-dependent activating DREADDs (referred to as Gq) in hypothalamic cells projecting to the ventral striatum (Fig. 4E). We unilaterally injected WT mice with Cre-dependent Gq DREADDs in the hypothalamus and AAV2-MNM004-Cre vector into the ventral striatum (Fig. 4F). As chemogenetic receptors lack biological ligands, activation depends on the presence of inert exogenous agonists (e.g., clozapine-N-oxide (CNO)). At 8 weeks post-injection, we administered CNO intraperitoneally and perfused the mice 2 hours later to assess the cell activity using immunohistochemistry. We observed an increase in the cFos cell-activation marker exclusively in the Gq-injected side of the hypothalamus, which demonstrates selective activation of hypothalamus neurons that project to the ventral striatum using this approach (Fig. 3G).

**Figure 4.**
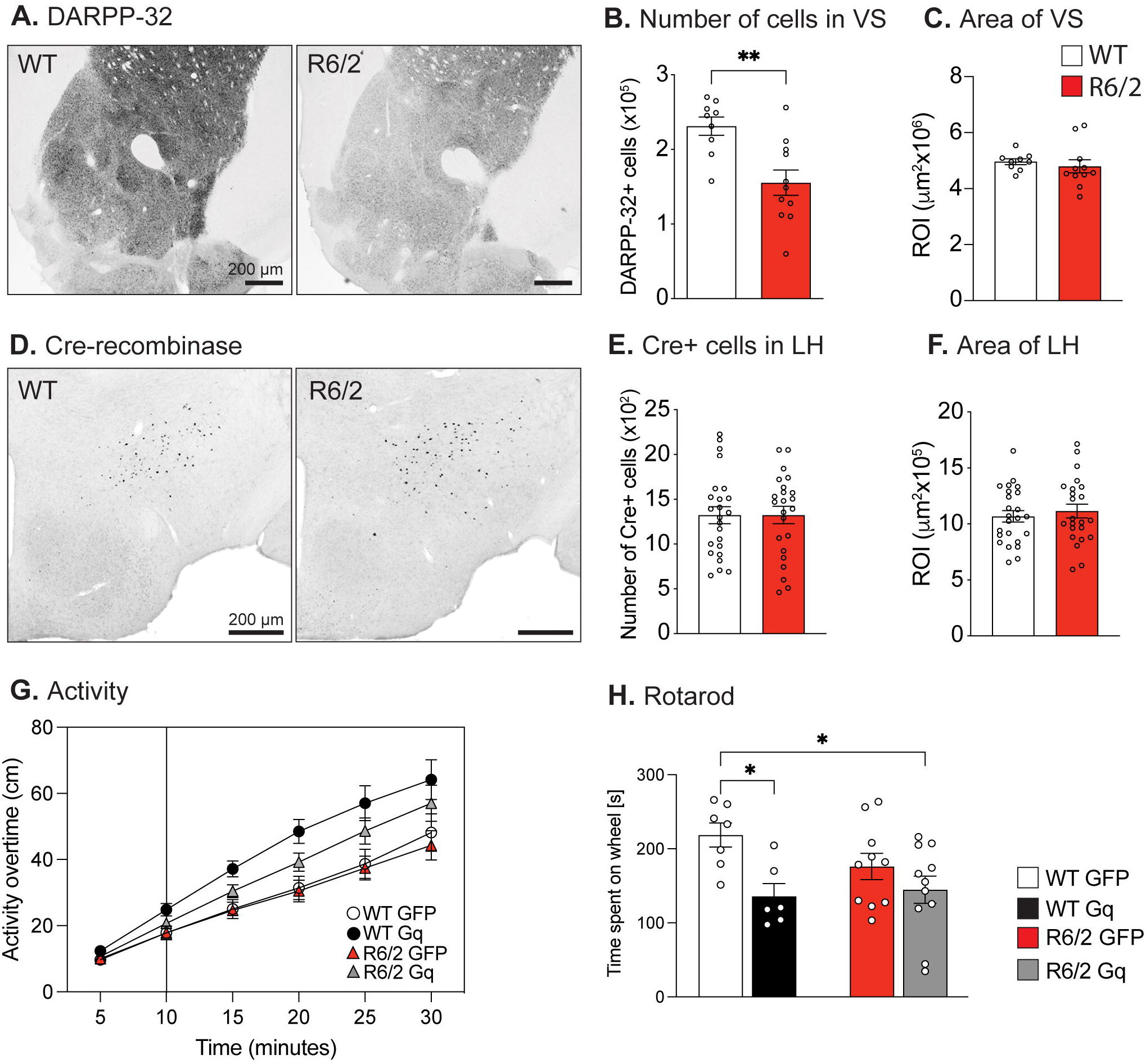
Assessment of ventral striatum neuropathology in R6/2 mice and chemogenetic activation of the neurocircuitry. **(A)** Representative images show DARPP-32 immunohistochemistry in the WT and R6/2 striatum. **(B)** The ventral striatum DARPP-32 positive cells stereological assessment shows a reduction in R6/2 mice compared to WT littermate controls (unpaired t-test, p=0.0026). **(C)** The ventral striatum area was not altered in R6/2 mice compared to WT littermate controls. **(D)** Cre-recombinase immunostaining showing the Cre positive cell bodies in WT and R6/2 mice. **(E)** Stereological assessment of Cre positive cells in WT and R6/2 mice hypothalamus at 8 weeks post-injection (unpaired t-test, p=0.99) **(F)** with comparable lateral hypothalamic area assessed for both genotypes. **(G)** General activity measurement over 30 minutes after i.p. CNO administration at time point zero (three-way repeated-measures ANOVA, the effect of time F (1.135, 32.91) = 263.2, p<0.0000001, the effect of vector F (1, 29) = 10.58, p=0.0029, the effect of genotype F (1, 29) = 1.521, p=0.227). **(H)** Rotarod behavior test showing time spent on rotating rod and reduced time on rod reveal a motor deficit (two-way ANOVA, effect of vector F (1, 30) = 9.159, p=0.005). Points on scatter graphs represent individual mice, the bars are means, and the whiskers indicate ± SEM.

### Chemogenetic activation of hypothalamic to the ventral striatum circuitry revealed a blunted response in R6/2 mice and deteriorated rotarod performance in WT mice

The R6/2 and the AAV vector-mediated animal models recapitulate clinical HD hypothalamic pathology with hypocretin cell loss [21, 23]. Therefore, to assess if the hypothalamus to ventral striatum neurocircuitry function is altered in HD, we used the R6/2 mouse model. First, we assessed whether the R6/2 mouse model exhibits ventral striatum neuropathology. We quantified the number of DARPP-32 cells in the ventral striatum. R6/2 mice had ∼33% reduction in DARPP-32 cells compared to WT littermates (Fig. 4A and 4B) without ventral striatum atrophy at 10 weeks of age (Fig. 4C). To examine whether retrograde transport of Cre was hampered in R6/2 mice due to hypothalamic neuropathology, we first injected R6/2 and WT mice unilaterally with AAV2-MNM004-Cre in the ventral striatum at 7 weeks. We quantified the Cre-expressing cell bodies in the lateral hypothalamus 6 weeks post-injection (Fig. 4D). The number of cells positive for Cre was comparable in R6/2 and WT mice, denoting that a similar number of projecting cells in the hypothalamus was targeted using this approach (Fig. 4E and 4F). We then explored the effects of activation of hypothalamic projection neurons in the R6/2 mouse model. To determine how stimulation of hypothalamic neurons affects behavior in R6/2 mice, we expressed Gq DREADDs selectively in hypothalamic-to-ventral striatum projecting neurons of R6/2 and WT mice. We employed a Cre-dependent rAAV8-mCherry-GFP flip switch vector as a virus control group (Fig. 3A–C). Both WT and R6/2 mice were injected at 9 weeks of age, and behavioral tests were performed at 5 weeks post-injection. Since CNO administration may exert off-target effects due to back-metabolization of clozapine and other clozapine metabolites [42], we injected all four groups of mice with the same dose of CNO to account for these potential off-target effects. First, we assessed general ambulatory movement using an open-field test, and we injected both mCherry and Gq groups intraperitoneally with CNO at time 0 and followed the activity. A repeated two-way ANOVA showed that there was a significant effect of time (p=0.0004), genotype (R6/2 vs. WT; p=0.0042), and vector (mCherry vs. Gq; p=0<0.0001) over the first 20 minutes (Fig. 4G). Even though activation led to increased activity, the rotarod performance of WT mice deteriorated as time spent on wheels decreased by ∼38%. The CNO administration had no effect on the rotarod performance of R6/2 mCherry groups compared to the R6/2 Gq group indicating an altered circuitry function in HD mice (Fig. 4H). We employed the limb clasping test to assess functional motor impairments, but activation of the hypothalamus to ventral striatum neurocircuitry had no effect (Supp. Fig. 4A).

Given the role of the ventral striatum in reward-related behavior and motivational force, the activation of hypothalamic-ventral striatal projections may affect psychiatric-like phenotypes such as anxiety, fear, and obsessive-like behavior [43]. As a result, these psychiatric traits could also affect motor performance. Therefore, we profiled mice using elevated plus maze and marble burying tests and assessed the time spent and distance moved in the center of the open-field arena. Since there were no effects on the anxiety-like, fear, or obsessive-like phenotype, we can conclude that psychiatric traits did not evoke or worsen motor activity (Supp. Fig. 4B and 4G). Lastly, we evaluated the reward-related behavior using a reward baited open-field test. However, there was no effect of vector or genotype (Supp. Fig. 4H and 4I). Taken together, activation of the hypothalamic-ventral striatal circuitry in R6/2 mice showed no beneficial effect, and the absence of response compared to WT mice suggests a dysfunctional circuitry in the R6/2 mice.

## Discussion

In the present study, we report for the first time that selective overexpression of mHTT in the hypothalamus can lead to neuropathology in the ventral striatum of mice. We show that neuropathology constitutes a neuronal loss, including loss of DARPP-32 immunopositive neurons, which are selectively affected in HD [44]. As mHTT is expressed only in the hypothalamus using regional targeting with AAV vectors in the present study, the neuropathology in the ventral striatum is mediated by non-cell autonomous effects of mHTT. Non-cell-autonomous effects of mHTT in the dorsal striatum have previously been attributed to the action of mHTT in the cerebral cortex and cortico-striatal pathways [6-9]. Although recent work has suggested that the synaptic disconnection generated by mHTT may extend to other excitatory pathways, the circuitry between the hypothalamus and the ventral striatum has been overlooked [45]. Our results show that non-cell-autonomous effects via the hypothalamus-ventral striatum pathway may also play a role in striatal neuropathology in HD. Furthermore, pathology of the ventral striatum is recapitulated in the R6/2 HD mouse model, where the hypothalamus-ventral striatum circuitry is disrupted as indicated by a blunted response using DREADD-mediated activation. Hence, the spectrum of HD pathology includes ventral striatal pathology via non-cell autonomous hypothalamic ventral striatal projections.

Projections from hypothalamic neurons to the ventral striatum are part of the reward circuitry [16, 38, 39]. In the present study, we show that a major part of the hypothalamic projections neurons to the ventral striatum are hypocretin immunopositive. This is in line with recent work showing direct excitatory hypocretin projections from the hypothalamus to the ventral striatum [15]. Hypocretin is a neuropeptide involved in regulating emotion, metabolism, and sleep and has also recently been implicated in the control of movement and cognition [46-48]. Hypocretin cell loss is found in both clinical HD and several HD mouse models [3, 20, 21, 49, 50]. Hypocretin neurons signal both via the neuropeptides hypocretin-A and hypocretin-B as well as via glutamate [15, 51, 52]. It is possible that hypocretin neuropathology mediates pathological effects in the ventral striatum via glutamatergic dysfunction. This may be similar to the proposed potential underlying mechanisms of cortico-striatal dysfunction mediated by an altered excitotoxicity [45]. Hence, pathology in the hypothalamus may play a role in contributing to ventral striatum pathology in HD via glutamatergic dysfunction.

The dorsal striatum pathology in HD is well characterized and is mainly linked to movement disorder in HD, while the ventral striatum has been less investigated. The early postmortem studies on clinical HD accumbens showed an increase in dopamine content and less shrinkage of the region compared to the dorsal part [53]; however, ultrastructural analysis of neuronal bodies using electron microscopy showed nuclear membrane indentation of more than 25% in HD accumbens as compared to less than 5%, and 2% in the frontal cortex and caudate nucleus, respectively [11]. In the original Vonsattel classification of neuropathology in HD, no macroscopic findings were observed in this area, even in otherwise severely affected cases (Vonsattel et al., 1995). With the increased awareness of non-motor aspects of HD, including psychiatric symptoms such as depression, anxiety, apathy, and altered social cognition, as well as metabolic dysfunction with increased appetite, more detailed studies on the ventral striatum have appeared. An early atrophy of the ventral striatum in clinical HD has been found in imaging studies [54], and postmortem human HD studies have identified loss of neurons in this area [55]. The present study shows that the commonly used R6/2 mouse model exhibits ventral neuropathology with a 33% loss of DARPP-32 immunopositive neurons. In our experimental paradigms using AAV vectors for targeted transgene expression, we show that ventral striatum neuropathology can be caused both by non-cell autonomous effects via hypothalamic projections and by direct expression of mHTT in this area. The consequences of pathology in the ventral striatum in HD are not fully known. In our models, we show that even up to 10 months post-injection of AAV vectors expressing mHTT in the ventral striatum, there is no effect on body weight. This contrasts with the effects of selective expression of mHTT in the hypothalamus, which leads to rapid body weight gain and severe metabolic imbalance in mice [21, 28]. These experiments show that it is the direct consequence of mHTT in the hypothalamus and not in the ventral striatum that leads to metabolic changes in mice. Furthermore, direct expression of mHTT in the ventral striatum did not lead to any effects on the motor or anxiety-like behavior at the time point studied in our experiments. Alterations in these behaviors could appear later, but it is also possible that the HD ventral striatum lies behind these behavior phenotypes concomitantly with hypothalamic pathology and alterations. In the present study, we did not test if the ventral striatal expression of mHTT led to depressive-like behavior. Other studies have found that the ventral striatum is involved in depressive-like behavior via cyclin-dependent kinase 5 in the Hdh^+/Q111^ knock-in mutant mice [56]. In other animal studies, motivational deficits have been linked to altered dopaminergic signaling in the ventral striatum of Q175 knock-in mice of HD [57], and reduced BDNF expression in this area has been linked to altered social interaction in the BACHD rat [58]. Hence, the ventral striatum may play a role in mediating several psychiatric symptoms of HD.

Chemogenetics has emerged as a useful method to investigate the functional role of specific neuronal circuitries for behavioral outcomes [26]. Since the discovery by the Roth laboratory [25], it has been used to shed light on the role of several neuronal circuitries in behavior, but criticism has also arisen mainly regarding the most used ligand CNO. There is a debate regarding whether CNO permeates the blood-brain barrier and whether it is its metabolite clozapine that activates the DREADD receptors. Clozapine is an approved antipsychotic drug that acts on many different neurotransmitter receptors, and such action may also lead to off-target effects [42]. Nevertheless, low concentrations of CNO in the order of magnitude used in this study cause sub-threshold levels of clozapine that are unlikely to act on other receptors than DREADDs due to their high-affinity [40, 42, 59]. Here we used conditional flip switch vectors expressing Gq DREADDs to selectively activate hypothalamic-to-ventral striatum projecting neurons of R6/2 and WT mice. To control for the possibility that CNO administration may exert off-target effects due to a back-metabolization to clozapine and other clozapine metabolites, we have injected all four groups of mice with the same dose of CNO [42]. We tested the mice using a battery of tests examining motor function as well as anxiety, reward, and obsessive-like behavior. Activation of this pathway appeared to increase general motor activity and worsen performance on the rotarod in wild-type mice. This may be linked to activating the recently identified lateral hypothalamico network of movement-promoting neurons regulated by hypocretin [46]. The response was blunted in R6/2 mice, suggesting that this circuitry is dysfunctional.

This study also revealed unfavorable effects of long-term wtHTT expression as a control for HTT CAG repeats and vectors. Previously, we showed that the expression of 853 aa long wtHTT fragments led to a milder degree of hypocretin cell loss than the mHTT group at one-year post-injection, 28% vs. 78% loss, respectively, whereas expression of GFP had no metabolic effect or hypocretin neuropathology [28], indicating that these effects are rather related to region-specific HTT functions and not due to any aberrant protein overexpression. In this study, the ventral striatal targeted overexpression of wtHTT and mHTT with 863 aa caused a similar degree of DARPP-32 cell loss in this area, and we show that 171 aa long wtHTT and mHTT hypothalamic overexpression led to a comparable hypocretin cell loss at 8 months post-injection. Furthermore, the degree of ventral striatal pathology aligned with hypothalamic neuropathology. Several studies showed that wtHTT overexpression alters gene expression and modifies the HD phenotype. In the dorsal striatum, expression of HTT171-18Q led to mild alterations than HTT171-82Q in the transcriptome profile at 6 weeks post-injection [60], and the hypothalamic overexpression of wtHTT led to major transcriptional alterations at 4 weeks post-injection [21, 29]. Apart from region-specific overexpression of wtHTT expression, HD transgenic mice expressing full-length wtHTT (i.e., YAC18, Hu18/18) also exhibited an altered metabolic phenotype [36, 61, 62]. In summary, studies set out to investigate mHTT disease mechanisms compared to wtHTT, as control groups, could lead to misrepresentations, as wtHTT expression may alter the phenotype in diverging or converging ways to mHTT.

In conclusion, our results indicate that expression of mHTT in the hypothalamus can lead to neuropathology with cell loss in the ventral striatum in mice. Furthermore, pathology of the ventral striatum is recapitulated in the R6/2 HD mouse model, where the hypothalamic-ventral striatal circuitry is disrupted as indicated by a blunted response using DREADD-mediated activation. Hence, ventral striatal pathology in HD may be mediated by non-cell autonomous effects of mHTT in hypothalamic-ventral striatal projections.

## Acknowledgments

We thank Ulla Samuelsson, Ulrika Sparrhult-Björk, Björn Anzelius, Anna Hansen and Anneli Josefsson for excellent technical assistance.

## Author contributions

RSK, ÅP, TB, and MB were responsible for the study conception and design. RSK, NA, MD, TB performed the experiments and analyzed the data. The first draft of the manuscript was written by RSK and ÅP. All authors read and approved the final manuscript.

## Declarations

### Disclosure of potential conflicts of interest

#### Funding

The research was supported by research grants to AP from the Swedish Research Council (grant numbers 2018/02559 and 2022/01092), the ALF system at Region SkÅne as well as the Knut and Alice Wallenberg Foundation (# 2019.0467), Swedish Brain Foundation research grant to AP, RSK, and MB (FO2017-0074) and The Royal Physiographic Society of Lund, Thorsten and Elsa Segerfalk Foundation, Neuroförbundet, Fredrik O Ingrid Thurings Foundations to RSK. RSK has been supported by the Swedish Brain Foundation (#PS2017-0057) and Swedish Society for Medical Research Postdoctoral Fellow grants (#P18-018).

### Conflict of Interest

The authors report no conflicts of interest.

### Ethics approval

The experimental procedures performed on mice were carried out in accordance with the approved guidelines in the ethical permit approved by the Lund University Animal Welfare and Ethics committee in the Lund-Malmö region (ethical permit numbers M20-11 and M65-13 and M15499-2017).

## Supplementary figures

**Supplementary Figure 1. Expression of short mHTT and wtHTT fragments selectively in the hypothalamus leads to ventral striatum cell loss. (A)** Immunohistochemical staining of DARPP-32 showing the striatum of uninjected mice and mice injected in the hypothalamus with AAV5-HTT171-18Q and AAV5-HTT171-79Q. **(B)** Stereological quantification of ventral striatum DARPP-32 positive cells shows a reduction in 171HTT-79Q and HTT171-18Q groups compared to uninjected controls (One-way ANOVA, p=0.0037). **(C)** NeuN immunolabeling and **(D)** quantification in the ventral striatum showed reduced ventral striatum cells in both 171HTT-79Q and HTT171-18Q groups (One-way ANOVA, p=0.0004). **(E, F, G)** The number of neurons and glia were assessed using Cresyl violet staining, and while the number of neurons was reduced in both HTT groups (One-way ANOVA, p<0.0002), the number of glial cells was not altered (Kruskal-Wallis test, p=0.56). **(H)** Hypocretin immunolabeling and **(I)** quantification of the number of hypocretin-positive cells showed a significant reduction in mHTT and wtHTT groups compared to uninjected controls (One-way ANOVA, p=0.0062). **(J)** Body weight changes over 32 weeks post-injection time (repeated measures two-way ANOVA, fitted Mixed-effects model (REML), n=13-14/group 0-4 weeks, and n=6-7/group between 4-16 weeks, n=7/group between 17-32 weeks, Time effect p <.001, F (1.450, 28.20) = 76.82, Vector p=017, F (2, 38) = 4.544). **(K)** GFP immunohistochemistry shows the absence of GFP-positive cells in the ventral striatum after 4 weeks of unilateral AAV5-Cre delivery in the hypothalamus of the R26R-EYFP reporter mouse. Points on scatter graphs represent individual mice, the bars are means, and the whiskers indicate ± SEM. Points on scatter graphs represent individual mice, the bars are means, and the whiskers indicate ± SEM.

**Supplementary Figure 2. Expression of mHTT fragments in the ventral striatum of mice does not affect body weight and behavior phenotype at 10 weeks post-injection. (A)** The photomicrographs of GFP and HTT transgenes in the ventral striatum. **(B)** Expression of HTT fragments in the hypothalamus did not affect body weight at 10 weeks post-injection (Kruskal-Wallis test, p=0.39). Assessment of anxiety-like phenotype showed no change in **(C)** activity (Kruskal-Wallis test, p=0.51), **(D)** time spent on open arms (One-way ANOVA, p=0.31), and **(E)** open arm entry frequencies (One-way ANOVA, p=0.58). **(E)** The rotarod performance (One-way ANOVA, p=0.21), **(F)** general activity (Kruskal-Wallis test, p=0.25), and **(G)** activity in the arena center (Kruskal-Wallis test, p=0.12) were also comparable. Points on scatter graphs represent individual mice, the bars are means, and the whiskers indicate ± SEM.

**Supplementary Figure 3. A large portion of hypothalamic projection neurons to the ventral striatum comprises lateral hypothalamus hypocretin cells. (A**) The maximum intensity projection and the orthogonal projections of Z-stack images illustrate the colocalization (cyan) of hypocretin-positive cells (blue) with projection cells (GFP transgene, green). The maximum intensity Z-projection images showing (mCherry, red) showed hypothalamic projection neurons were not **(B)** oxytocin, **(C)** vasopressin, or **(D)** TH positive cells in zona incerta area (green) in WT mice expressing Gq DREADDs selectively in hypothalamic-to-ventral striatum projecting neurons.

**Supplementary Figure 4. The hypothalamus to ventral striatum neurocircuitry profiling in R6/2 and WT mice using chemogenetic activation. (A)** Hindlimb clasping functional motor test shows a genotype effect (two-way ANOVA, effect of vector F (1, 29) = 8.043, p=0.0082). **(B)** Elevated plus maze test showing no change in total distance moved. **(C)** There is an effect of vector and genotype for open arm entry frequency (two-way ANOVA, the effect of genotype F (1, 29) = 15.65, p=0.0005; effect of vector F (1, 29) = 5.947, p=0.0211), the effect of genotype x vector F (1, 29) = 6.262, p=0.0182)), and **(D)** genotype effect for the duration in open arms (two-way ANOVA, effect of vector F (1, 29) = 13.83, p=0.0009). **(E)** Open field arena entrance frequency showed no effect of vector or genotype. **(F)** Time spent in the center of the open field arena showed no effect of the vector (two-way ANOVA, effect of vector F (1, 29) = 0.3928, p=0.54, effect of genotype F (1, 29) = 8.844, p=0.006)). **(G)** There was no significant effect of genotype or vector in the marble burry test. **(H)** Locomotor activity in the reward-baited arena over 10 minutes and **(I)** consumed peanut butter were comparable. Points on scatter graphs represent individual mice, the bars are means, and the whiskers indicate ± SEM.

## Material and methods

### Animals

The experimental procedures were performed on mice following the guidelines in the ethical permits approved by the Lund University Animal Welfare and Ethics committee in the Lund-Malmö region (ethical permit numbers M20-11, M65-13 and M15499-17). All mice were housed with free access to water and a standard chow diet in an animal facility with 12 hours a night/day cycle.

The experiments using stereotactic injections of AAV vectors expressing huntingtin fragments, or green fluorescent protein (GFP) in the hypothalamus or the ventral striatum were carried out in female mice of the FVB/N strain at two months of age and weighing 20-26 g (The Jackson Laboratories) and hypothalamic Cre-recombinase expression experiments were carried out on R26R-EYFP mice (B6.129×1Gt(ROSA)26Sortm1(eYFP)Cos/J, C57BL/6 strain; Jackson Laboratories). The female R6/2 mice and their WT littermates from B6/CBA strain were utilized at 10 weeks of age for neuropathological analysis of the ventral striatum and a 6-7 weeks old group for chemogenetics modulation of the hypothalamus to ventral striatum neurocircuitry. The R6/2 mice were generated by crossing heterozygous R6/2 males with WT females. The genotyping was performed as described previously [22] and the R6/2 mice used in this study carried a CAG repeat size between 273-285, resulting in a slower disease progression compared to the original R6/2 line with 150 CAG repeats [63].

### Adeno-associated viral vectors

We employed a number of recombinant adeno-associated viral (rAAV) vectors in this study. *HTT and GFP expressing AAV vectors:* To express different fragments and CAG repeat size of HTT proteins in the hypothalamus and ventral striatum, we employed AAVs carrying a mutant (79Q, polyglutamine) or a wild-type HTT (18Q). Each HTT vector contained either 853 (long fragment) or 171 (short fragment) amino acid length N-terminal region of the human HTT gene [21]. The HTT vectors were expressed under the human Synapsin-1 (hSyn) promoter, and the wild-type HTT carried 18 CAG repeats (HTT853-18Q or 171HTT-18Q), whereas the mutant form carried 79 CAG repeats (HTT853-79Q or 171HTT-79Q). The viral vectors were pseudotyped rAAV2/5 vectors (referred to as AAV5 elsewhere). The AAV5 vectors expressing GFP under the Syn-1 promoter constituted a control group for the ventral striatum 10 weeks post-injection. The AAV vectors were obtained with a double-transfection method utilizing the helper plasmid encoding essential adenoviral packaging genes, as described before [64].

#### Chemogenetics and retrogradely transportable vectors

The activating double floxed Gq-coupled hM3D DREADD construct was fused with mCherry under the control of the human syn promoter, and it was packaged into an AAV8 capsid (pAAV-hSyn-DIO-hM3D(Gq)-mCherry, RRID: Addgene_44361) [65]. The retrogradely transportable AAV2-MNM004-Cre and AAV8-mCherry-CTE-GFP AAV vectors were produced using PEG precipitation and chloroform extraction as previously described [66]. In brief, the self-complementary CMV-Cre genome was packaged into MNM004 [27], and the color-changing vector hSyn-mCherry/CTE/GFP was packaged into AAV8 using PEI transfection. HEK293 cells were triple transfected with ITR-flanked genomes. AAV capsid and the helper plasmid pHGT1. AAVs were harvested using polyethylene glycol 8000 (PEG8000) precipitation and chloroform extraction, followed by PBS exchange in concentration columns at 72-hour post-transfection. Purified AAV vectors were titered using droplet digital PCR [67] with primers specific for the ITRs (forward primer 5′-CGG CCT CAG TGA GCGA-3′ and reverse primer 5′-GGA ACC CCT AGT GAT GGA GTT-3′).

### Viral vector injections

Mice were subjected to 2% isoflurane in oxygen/nitrous oxide (3:7) anesthesia and then transferred onto a stereotactic instrument. The skull of the mouse was thinned with a dental drill at the chosen location, and the thin bone flap was carefully removed, leaving the dura intact. The stereotaxic coordinates for the hypothalamus and ventral striatal regions were determined according to bregma using the 3rd edition Franklin, and Paxinos brain atlas and dorsal-ventral coordinates were calculated from dura mater [68]. The hypothalamic coordinates were anterior-posterior (AP) = 0.6 mm, medial-lateral (ML) = 0.6 mm, dorsal-ventral (DV) = 5.3 mm and ventral striatum coordinates were AP = 1.6 mm, ML = 1.3 mm, DV = 4.0 mm. A pulled glass capillary with an outer tip diameter of ≈ 80 μm attached to a 5 μl Hamilton syringe (Nevada, USA) was used to inject the total viral vector volume of 0.5 μl for hypothalamic and 0.3μl for ventral striatum regions. The rate and volume of viral vector delivery were 0.05 μl injections in 15 s intervals, subsequently to an initial injection of 0.1 μl of viral vector solution. Following the injection, the glass capillary was left in the target for additional 5 minutes to allow absorbance of the virus by the tissue. The vector concentrations were as follows: The vectors and titers used for in vivo experiments were rAAV5-hSyn-HTT853-18Q (1.4E+14 GC/ml), rAAV5-hSyn-HTT853-79Q (1.6Ex14 GC/ml), rAAV5-hSyn-GFP (5.7E+13 GC/ml), rAAV5-hSyn-Cre (1.5E+14 GC/ml), rAAV5-hSyn-HTT171-18Q (1,2E+15 GC/ml), rAAV5-hSyn-HTT171-79Q (6,1E+14 GC/ml), AAV2-MNM004-CMV-Cre (2.9E+12 GC/ml) and AAV8-hSyn-mCherry-CTE-GFP (9.4E+12 GC/ml), and pAAV-hSyn-DIO-hM3D(Gq)-mCherry (9.7 E+12 GC/ml).

### Behavioral tests

Mice were transported in their home cages to the behavioral room and were allowed to acclimate to the room at least the day before testing. The behavioral tests for the chemogenetically modified mice were performed using clozapine nitric oxide (CNO, Sigma, 4 mg/kg). Both Gq and GFP-expressing mice were intraperitoneally injected with the same dose of CNO to account for CNO off-target effects. The behavior tests were performed following the CNO injections. The mice were placed on the open field arena right after the intraperitoneal CNO injections to assess the activity change in response to chemogenetic receptor activation, while the rotarod, hindlimb clasping, marble-burying, elevated-plus maze, and reward-baited open-field test were done at least 20 minutes after the CNO injections. The open field, clasping, and accelerating rotarod were performed in this order on the same day with 20 minutes rest between the tests at 7 weeks post stereotaxic virus injections. The animals were rested for 1 week, and then at week 8 post stereotaxic injections, all the mice were subjected to first marble bury, then elevated-plus maze on the same day 30 minutes after the CNO injections. Lastly, at 9 weeks post stereotaxic virus injections, the mice were tested on reward-baited open field test 20 minutes after the CNO injections.

### Open Field Test

General locomotor activity was assessed using the open field test (OFT). Animals were placed in the center of the activity box (40 cm × 40 cm) enclosed by opaque walls. The movement of the animals across the arena was recorded and tracked with the AnyMaze behavior tracking software (Version 4.99M, Stoelting Europe, Dublin, Ireland, UK). The locomotor activity for the ventral striatum expression of HTT vectors was recorded for 1 hour, and for the chemogenetics experiments, mice were tracked for 2 hours immediately following the intraperitoneal CNO injections. The analysis of the recordings to calculate the total and overtime activity, time spent in the center zone and center zone entry frequencies was done using the Ethovision XT software (Version 13.0.1216, Noldus Information Technology, Wageningen, Netherlands).

### Accelerating rotarod test

To assess motor coordination and balance [69] an accelerating rotarod test was used (Rota-rod/RS, Panlab Harvard Apparatus). The rod diameter was 3 cm, and the fall height was 20 cm. The mice were trained on a fixed speed (2 minutes at 4 rpm) rotating rod for 3 consecutive days, and on the 4th day, mice were tested three times with a linear accelerating speed (5 minutes from 4 - 40 rpm). The mice were allowed to rest 30 minutes between each session, and the rotarod apparatus was cleaned with 75% ethanol between each trial. The latency to fall was recorded, and the data were expressed as the mean of three trials.

### Hindlimb clasping

Hindlimb-clasping phenotypes have been described in HD animal models. The hindlimb-clasping test relies on when a healthy WT mouse is suspended by the tail towards a horizontal surface. Healthy adult rodents splay all four limbs outwards, away from the abdomen [70]. Mice were lifted by the base of the tail from a horizontally placed metal grid at a distance just within reach for the forelimbs. The mice were suspended for 30 sec while recorded with a video camera. By a blinded rater to genotype and vectors, the hind-clasping behavior was scored from 0 to 3 according to criteria by Guyenet et al. [71], where a score of 0 represented no motor phenotype.

### Marble burying test

The marble-burying and digging behavior is a sign of anxiety and repetitive behavior in rodents [72]. For the marble burying test, 12 glass marbles in 1.5 cm diameter were equally spaced on the surface of the bedding of a clean home cage (40 cm length × 25 cm width × 20 cm height) containing approximately 4 cm high bedding material. The mice were kept in the cage for 30 min, and at the end of the session, the number of marbles buried was counted.

### Elevated-plus maze

The anxiety-like behavior was assessed using an elevated plus maze (EPM). The EPM consisted of two enclosed and two open arms (platform: 30 cm long and 6 cm wide and walls: 30 cm), and the platform was elevated 50 cm above its base. Mice were placed in the center of the maze, and the movement of the animals across the arena was recorded and tracked with the AnyMaze behavior tracking software for 5 minutes (Version 4.99M, Stoelting Europe, Dublin, Ireland, UK). The time spent on the open arms, open arm entry frequency and general movement, were assessed using the Ethovision Software (Version 3.1, Noldus Information Technology).

### Reward-baited open field test

The reward-baited open field test was adapted from Lockie et al. [73]. The mice were introduced to peanut butter on day 1. The peanut butter (PB) was provided in the home cage for 2 hours in a container secured to the cage bottom with Blu-Tac. On day 2, mice were food-restricted for 6 hours prior to the test. The PB container was secured to the center of a large open field arena surface (80 cm × 80 cm) with a 40 cm distance to each wall. The mice were allowed to explore the arena, and the activity was recorded and tracked with the AnyMaze behavior tracking software for 10 minutes (Version 4.99M, Stoelting Europe, Dublin, Ireland, UK). All the PB containers were weighed before and after each test. After each test, the arena was cleaned with 75% EtOH, and PB containers were replaced with new containers for each mouse.

### Immunohistochemistry

The mice were anesthetized with a terminal dose of sodium pentobarbital. The perfusion was performed transcardially first with saline and subsequently with pre-cooled 4% paraformaldehyde (PFA) at a 10 ml/min rate for 10 minutes. The brains were postfixed in 4% PFA solution at 4°C for 24 hours, then transferred to 25% sucrose solution at 4°C for ∼24 hours. For the immunohistochemical analysis, the brains were cut at 30 μm thick coronal sections in six series using a Microm HM450 microtome (Thermo Scientific). The brain sections were stored at −20 °C in an antifreeze solution containing 30% glycerol and 30% ethylene glycol in Phosphate-buffered saline (PBS) until further processing. Free-floating brain sections were rinsed with 0.05 M Tris-buffered saline (TBS) to remove the antifreeze solution 3 times for 10 minutes. An antigen retrieval step using 9.0 pH Tris/EDTA buffer solution in an incubator for 30 minutes at 80 °C is done for c-Fos immunohistochemistry. Next, the quenching reaction was performed in TBS with 3% H_2_O_2_ and 10% Methanol. The sections were pre-incubated for 1 hour at room temperature (RT) with blocking solutions containing 5% normal serum matched to the corresponding secondary and 0.25% triton-X in TBS (TBS-T). Next, the sections were left overnight at RT in primary antibody solution in TBS-T containing 5% normal goat serum. The primary antibodies dilutions were: c-Fos (1:1000, rabbit, #2250 Cell Signaling Technologies), anti-mCherry (1:5000, rabbit, #600-401-P16, Rockland), anti-tyrosine hydroxylase (1:2000; rabbit; Pel-Freez), anti-huntingtin (sc-8767; 1:500; goat; Santa Cruz), anti-ubiquitin (1:2000; rabbit; Dako), anti-GFP (ab290; 1:30000; rabbit; Abcam), vasopressin (1:10000, Chemicon) and oxytocin (1:2000, Phoenix Pharmaceuticals), anti-hypocretin (1:4000; rabbit; Phoenix Pharmaceuticals). Following a rinsing step (3 times for 10 minutes in TBS-T), the brain sections were incubated with secondary antibodies for 1 hour in 1% bovine serum albumin (BSA) containing TBS-T (1: 200; Vector Laboratories Inc.). Following the washing step with TBS-T, the sections were incubated with an avidin-biotin-peroxidase complex solution for 1h. The sections were washed with TBS-T, and then the staining was visualized using 3,3’-diaminobenzidine (DAB) and 0.01% H2O2 according to the manufacturer’s instructions. At last, the sections were washed with TBS and mounted on chromatin-gelatin-coated glass slides. For the Nissl stain cresyl violet (ICN Biomedicals Inc) staining, one series of sections were first mounted and then processed for staining according to the manufacturer’s instructions. For cover-slipping, the glass slides were first immersed in dH_2_O for 1 minute, then in increasing alcohol solutions (70%, 95%, 100%), cleared in xylene. The slides were secured with glass coverslips using Depex mounting medium (Sigma-Aldrich).

### Stereological analysis

To estimate the number of cells, unbiased stereological quantification principles were implemented [74]. Using Nikon Eclipse 80i upright microscope with an X–Y motorized stage (Märzhauser, Wetzlar, Germany); and a Z-axis motor with a high precision linear encoder (Heidenhain, Traunreut, Germany), first, the region of interest were delineated under the 4X objective and then the counting was performed using a 60X objective (Plan-Apochromat 1.4 N.A. oil immersion objective). The counting was performed with a random start systematic sampling routine (NewCast Module; VIS software; Visiopharm A/S, Horsholm, Denmark). All three axes and the input from the digital camera were controlled by a computer. The sampling interval was adjusted to count at least 100 cells for each brain.

### Confocal imaging

For the confocal imaging, following the washing step from the anti-freeze solution, brain sections were preincubated with 5% normal serum in PBS with 0.05% Triton-X-100. Next, the sections were incubated overnight at RT in 1% BSA in PBS with 0.05% Triton-X-100 solution containing primary antibodies. Next, the sections were incubated with primary antibodies overnight at room temperature. The primary antibody dilutions were vasopressin (1:10000, Chemicon) and oxytocin (1:2000, Phoenix Pharmaceuticals), anti-hypocretin (1:4000; rabbit; Phoenix Pharmaceuticals), anti-tyrosine hydroxylase (ab1542, 1:500; sheep; Chemicon) and anti-GFP (ab13970; 1:10000; chicken; Abcam). The sections were rinsed in TBS-T, then they were incubated with DyLight 488 (1:200; donkey anti-chicken, Jackson ImmunoResearch) and Cy5 (1:200; donkey anti-sheep; Jackson ImmunoResearch) secondary antibodies were diluted in TBS-T at RT for 1 hour. Lastly, the sections were washed two times for 10 minutes with TBS-T and once with TBS and then secured with coverslips using an anti-fading polyvinyl alcohol mounting medium for immunofluorescence imaging (PVA-DABCO, Sigma-Aldrich).

The sections were imaged using the Nikon Eclipse Ti-E inverted laser scanning microscope (Nikon, Instruments Inc., Melville, NY). The images were collected as Z-stacks, and the data was acquired in single-channel mode using Nikon EZ-C1 imaging software (v. 3.90), then imported to ImageJ (v. 1.48u4; NIH) and presented as orthogonal projection images.

### Gene expression analysis

Gene expression analysis of HTT was performed on snap-frozen hypothalamic and ventral striatum brain samples using liquid nitrogen, and the samples were kept at −80°C for further processing. The RNeasy Lipid Tissue Kit (Qiagen) was used for total RNA isolation. 1 μg of RNA samples were used for the reverse transcriptase reaction (SuperScript III Reverse Transcriptase kit, Invitrogen) according to the manufacturer’s instructions. The SYBR Green-based assay (SYBR Green I Master, Roche) with a two-step cycling protocol was used for the gene expression analysis using LightCycler 480 (Roche). Using the comparative ΔCt method (ΔΔCT method), the expression HTT gene was quantified relative to glyceraldehyde 3-phosphate dehydrogenase (GAPDH) and β-actin housekeeping genes. The primer sequences were as follows: HTT Forward 5’
s-AGTCAGATGTCAGGATGG-3’, Reverse 5’-CTGTAACCTTGGAAGATTAGAA-3’, β-actin: Forward −5’-GCTGTGCTATGTTGCTCTA-3’, Reverse 5’-TCGTTGCCAATAGTGATGA-3’; GAPDH: Forward 5’-AACCTGCCAAGTATGATGA-3’, Reverse 5’-GGAGTTGCTGTTGAAGTC-3’.

### Statistical analysis

The statistical analysis was done using Prism 9 software (Version 9.5.0, GraphPad). The data were tested for normal distribution using a Kolmogorov–Smirnov test and subsequently with Kruskal–Wallis followed by Dunn’s multiple-comparison test; one- or two-way ANOVA followed by Tukey’s multiple-comparison test; or unpaired t-test using. The three-way ANOVA was employed to analyze the relationship between three independent variables. The type of test used for each respective experiment is stated in the figure legends and results section. Data are presented as scatter plots with bars representing mean and whiskers ±SEM. The asterisks demark the p-value significance degree according to GraphPad style, as *P ≤ 0.05, **P ≤ 0.01, *** P ≤ 0.001, **** P ≤ 0.0001, and P > 0.05 considered nonsignificant.

## Notes

### Competing Interest Statement

The authors have declared no competing interest.

