## Supplementary figures and images for "Mutant huntingtin expression in the hypothalamus promotes ventral striatal neuropathology"

### Supplemental Figure 1

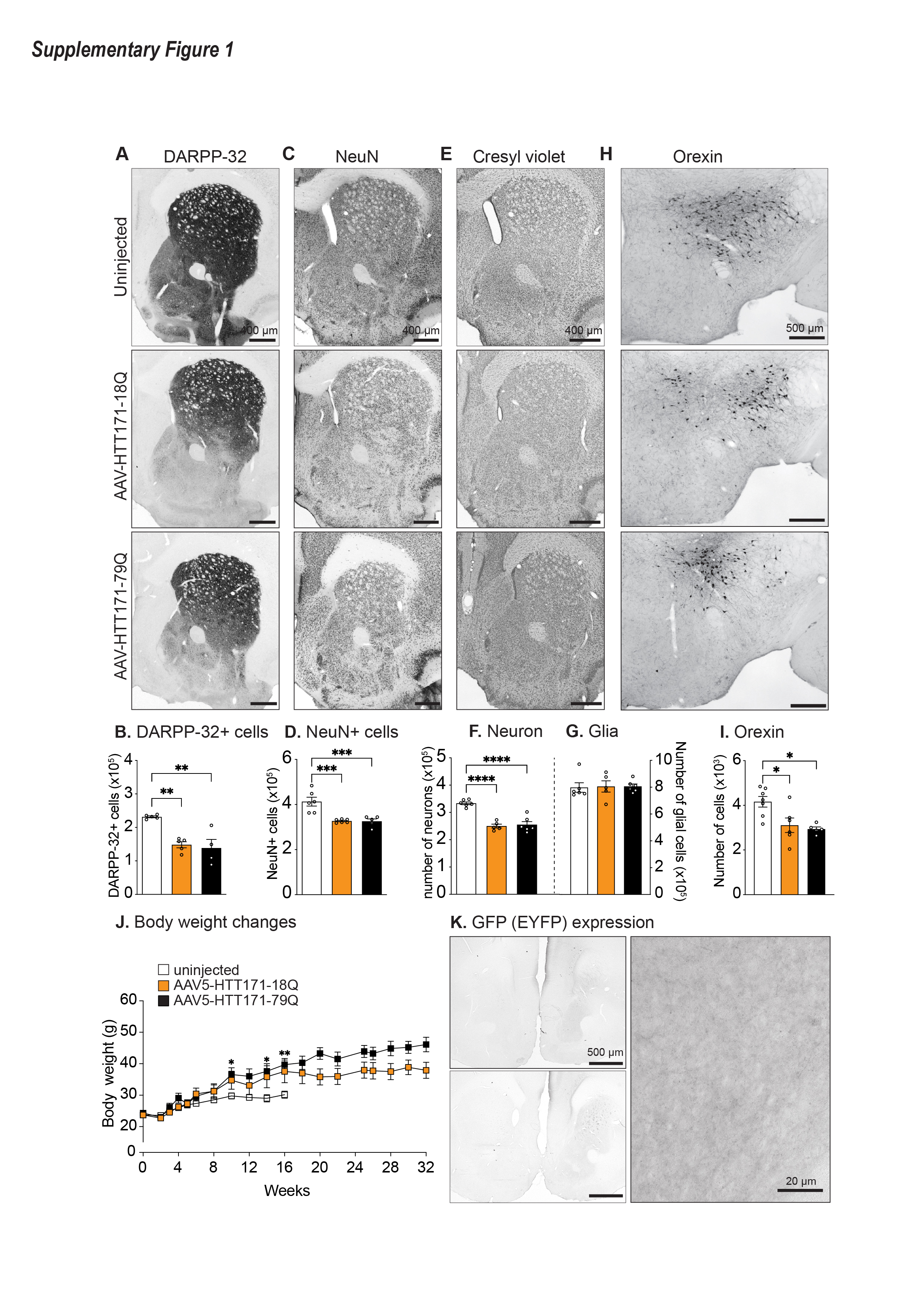

### Supplemental Figure 2

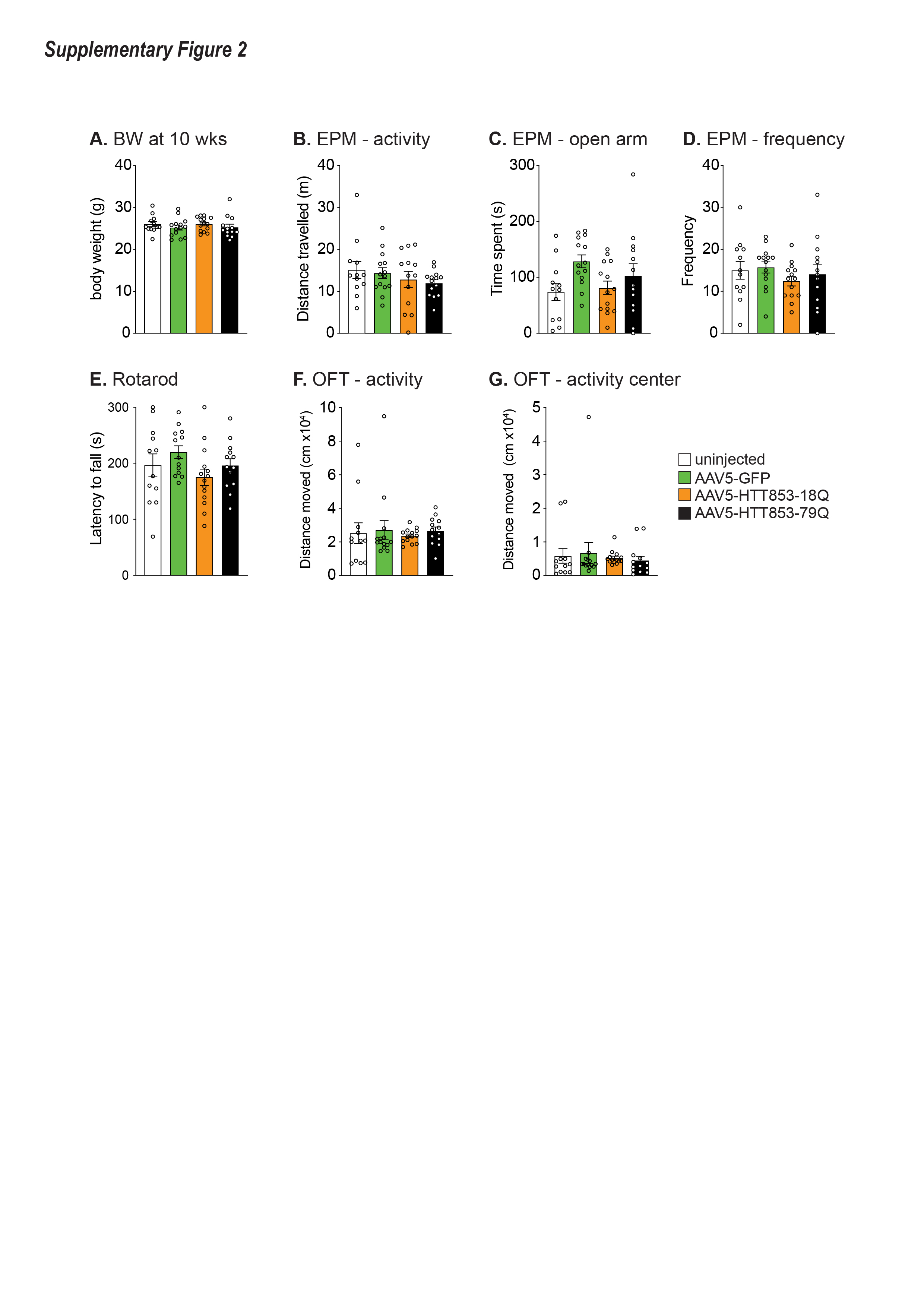

### Supplemental Figure 3

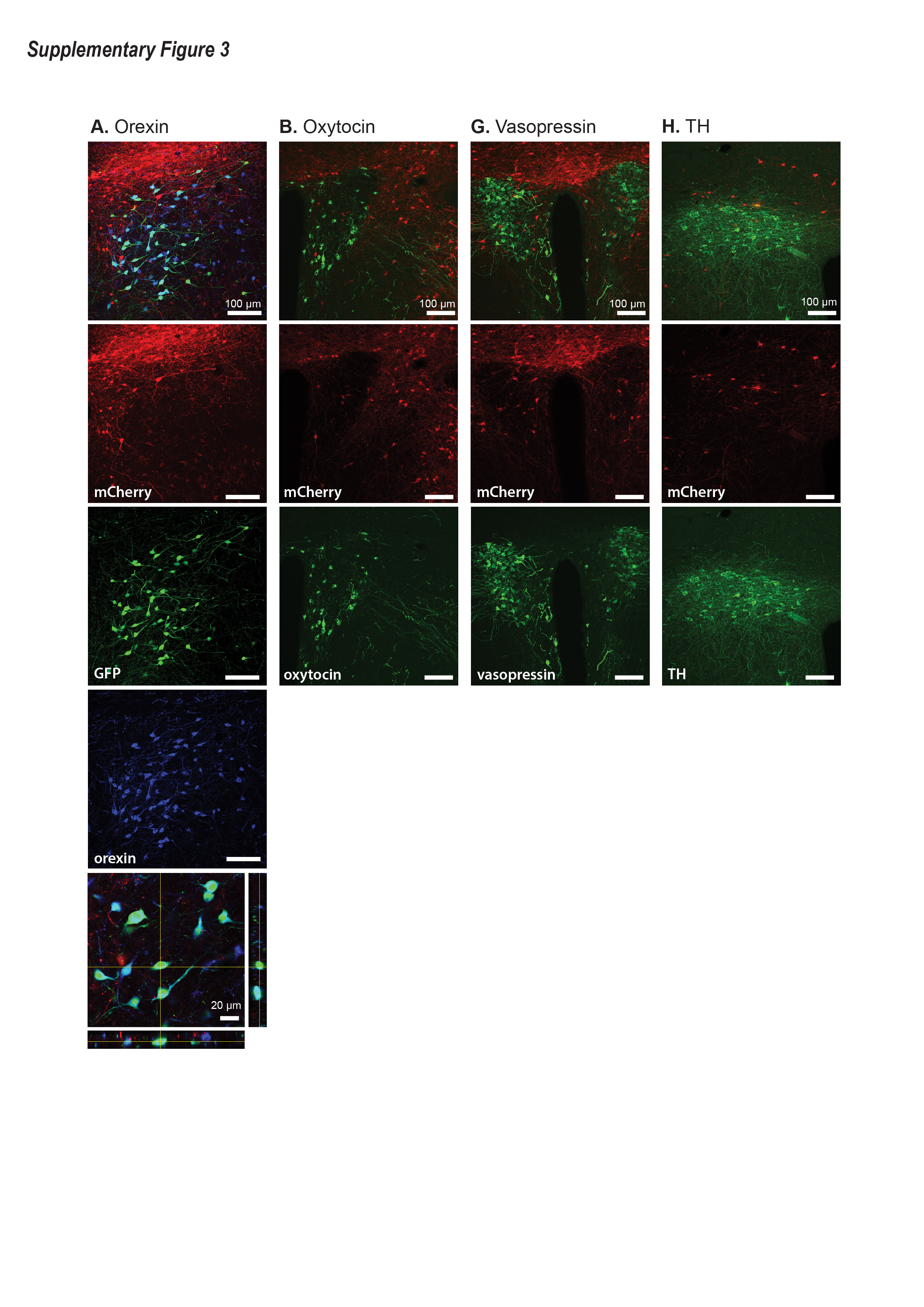

### Supplemental Figure 4

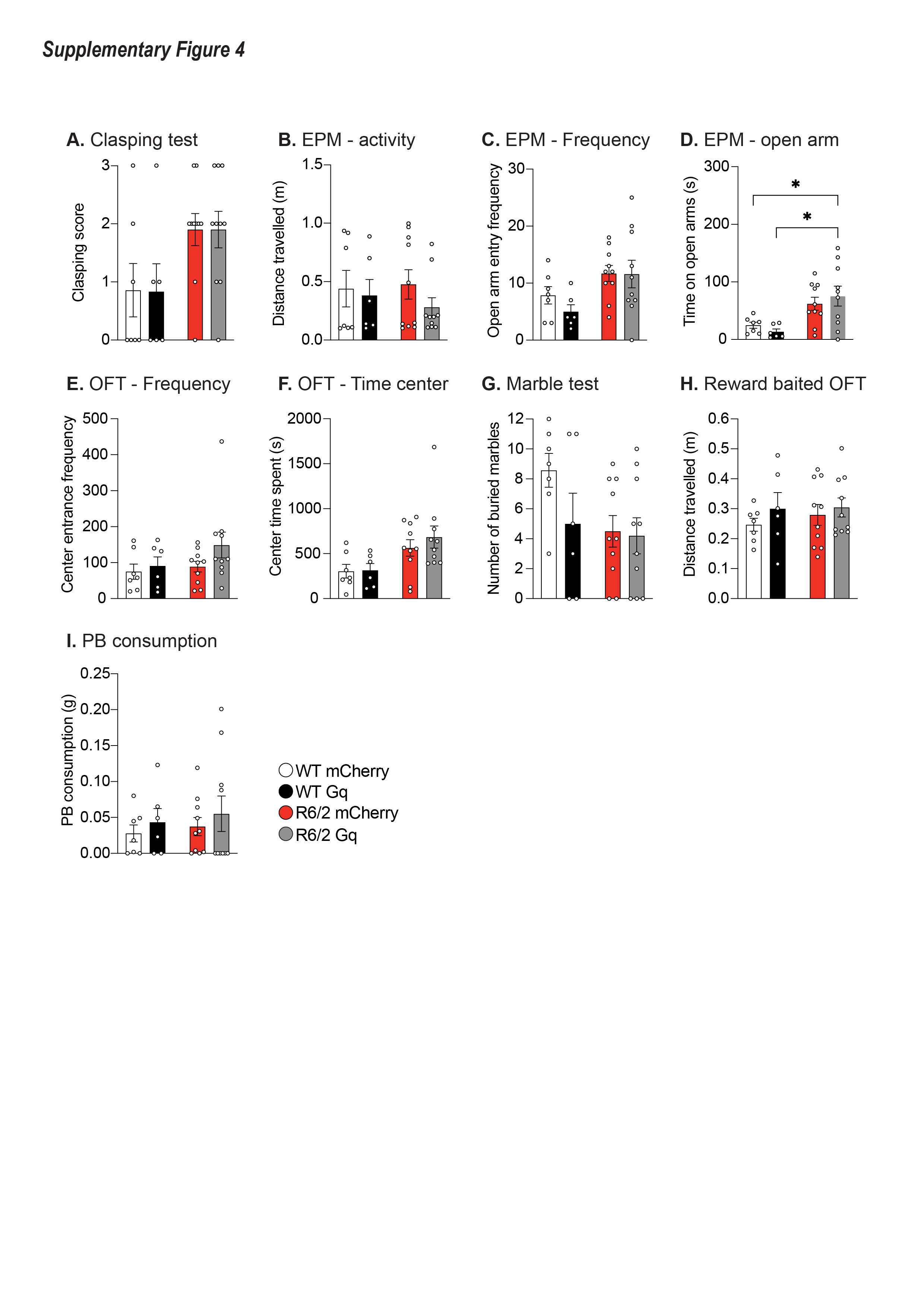
